# Homologous recombination deficiency in cell line libraries does not correlate with in vitro sensitivity to platinum agents or PARP inhibitors

**DOI:** 10.1101/2023.07.06.547853

**Authors:** Shiro Takamatsu, Kosuke Murakami, Noriomi Matsumura

## Abstract

Although preclinical studies for drug discovery and biomarker development are extensively conducted using large publicly available cancer cell line databases, there have been no reports to date that clarify the association between homologous recombination repair deficiency (HRD) and drug sensitivity using these data. We comprehensively analyzed the molecular profiles and drug response screening data from the Cancer Cell Line Encyclopedia. Unexpectedly, gene alterations in *BRCA1/2* or homologous recombination repair-related genes, HRD score, or mutational signature 3 were not significantly correlated with sensitivity to platinum agents or PARP inhibitors. Rather, higher HRD scores and mutational signature 3 were significantly associated with resistance to platinum and PARP inhibitors in multiple assays. These findings were consistent when analyses were restricted to breast and ovarian cancer cell lines and when data from the COSMIC Cell Line Project dataset were used.

**Significance:** In existing cancer cell line databases, the association between HRD status and sensitivity to platinum or PARP inhibitors differs from that expected from clinical tumor data. This discrepancy may also apply to other tumor characteristics, and researchers should be aware of the potential limitations of cell line data.

**Structural abstract:** *Background:* Comprehensive molecular profiling and drug sensitivity screening data from over 1000 cancer cell lines are currently available for preclinical studies including targeted drug discovery and biomarker development. However, there are no reports using these cell line databases to clarify the association between homologous recombination repair deficiency (HRD) and drug sensitivity.

*Methods:* We investigated the association between HRD status, including gene alterations in the homologous recombination repair (HR) pathway, HRD score, and mutational signature 3, and sensitivity to platinum agents and PARP inhibitors in the Cancer Cell Line Encyclopedia (CCLE) and the COSMIC Cell Line Project (CLP) datasets.

*Results:* In the CCLE dataset (n=1182), samples with *BRCA* alterations, including *BRCA1* methylation and *BRCA1/2* mutations with locus-specific loss-of-heterozygosity, exhibited higher HRD scores and mutational signature 3. These two scores were positively correlated (r=0.475, p=1.25 ×10^−52^). Unexpectedly, neither *BRCA1/2* nor HR-related gene alterations correlated with sensitivity to platinum agents or PARP inhibitors. Instead, significantly positive correlations were observed between drug-response AUC values and HRD scores in 60% (6/10) of platinum agent assays, and in 43% (6/14) PARP inhibitor assays, while no significant negative correlation was observed. Similar results were obtained in the analysis with mutational signature 3. These findings were consistent in analyses limited to ovarian and breast cancer cell lines and in the CLP dataset, indicating samples with HRD showed resistance rather than sensitivity to these drugs.

*Conclusion:* In existing cancer cell line databases, the association between HRD status and sensitivity to platinum and PARP inhibitors differs from that expected from clinical tumor data. This discrepancy may also apply to other tumor characteristics, and investigators should be aware of the potential limitations of cell line data.

## Introduction

Human tumor-derived models play a crucial role in cancer research and drug development. Among these models, human cancer cell lines have been extensively utilized in basic and preclinical research due to their ease of handling and low-cost availability. It has been reported that cancer cell lines robustly retain the genomic features of their original tumors (#1). The advent of next-generation sequencing has facilitated the comprehensive molecular profiling of more than 1000 representative cancer cell lines, and these datasets are available to researchers as public resources, including the Broad institute’s Cancer Cell Line Encyclopedia (CCLE) (#1) and the Sanger institute’s COSMIC Cell Line Project (CLP) (#2). In addition, in the past decade, several high-throughput drug screening studies using these cell lines have been conducted by different groups, including the Genomics of Drug Sensitivity in Cancer (GDSC) (#3), the Cancer Therapeutic Response Portal (CTRP) (#4, #5), and Profiling Relative Inhibition Simultaneously in Mixtures (PRISM) (#6), Genentech Cell Line Screening Initiative (gCSI) (#7), and others. Given the open availability of these datasets and their capacity to link specific molecular features to drug sensitivities, computational and machine learning-based approaches have been actively explored in recent years for targeted drug discovery and therapeutic biomarker identification (#1, #2, #8, #9, #10).

The homologous recombination repair (HR) pathway is the most precise and essential DNA damage repair mechanism for DNA double-strand breaks. Dysfunction of this pathway, known as homologous recombination deficiency (HRD), often caused by inactivation of BRCA1/2 molecules, leads to DNA damage accumulation and genomic instability that contribute to tumor development and progression. Cancer cells with HRD exhibit characteristic genomic changes referred to as the “Genomic Scar” signatures (#11, #12, #13). In ovarian and breast cancer, *BRCA1/2* mutations and genomic scar signatures are significantly associated with the sensitivity to PARP inhibitors and platinum agents (#14). Recently, their clinical utility as biomarkers has been investigated in other cancer types (#15, #16, #17).

Although various research has been conducted using the above cancer cell line databases, there have been no reports to clarify the association between HRD status and the sensitivity to PARP inhibitors or platinum drugs. Here, we examine the association between HRD status and in vitro drug sensitivity to these agents in the CCLE and the CLP datasets, demonstrating significant differences from that in clinical tumors. These findings highlight the limitations of drug and biomarker discovery research based on existing cell line databases and emphasize the need for caution when interpreting results obtained from such sources.

## Materials and Methods

### Cancer cell line datasets

The gene mutations, copy number aberrations, gene expression, and Reduced Representation Bisulfite Sequencing (RRBS)-based DNA methylation profiles for the Cancer Cell Line Encyclopedia (CCLE) dataset were obtained from the Dependency Map (DepMap) portal (Supplement TableX1).

For the COSMIC-Cell Line Project (CLP) dataset, the gene mutations, copy number aberrations, gene expression, and aneuploidy data were downloaded from the COSMIC website (GRCh37, release v97, 29th November 2022). In addition, the DNA methylation array data were obtained from the Gene Expression Omnibus (GEO) database (GSE68379). (Supplementary TableX1).

### HR-related gene mutations with locus-specific LOH

Based on a previous report (#18), 29 genes were defined as HR-related genes: *ATM, ATR, BARD1, BLM, BRCA1, BRCA2, BRIP1, CDK12, CHEK1, CHEK2, FANCA, FANCC, FANCD2, FANCE, FANCF, FANCI, FANCL, FANCM, MRE11, NBN, PALB2, RAD50, RAD51, RAD51B, RAD51C, RAD51D, RAD52, RAD54L*, and *RPA1*.

From the CCLE gene mutation profiles, mutations with ‘‘Variant_annotation’’ of ‘‘damaging’’ were obtained. From the CLP mutation profile, truncating mutations and missense mutations that were annotated as “deleterious” or “damaging” in all of 10 functional impact prediction methods (SIFT, Polyphen2, LRT, MutationTaster, MutationAssessor, FATHMM, PROVEAN, MetaSVM, MetaLR, M-CAP) were extracted.

As described in a previous report (#16), the estimated copy number of the minor allele at the locus of each mutation was evaluated. A value of 0 indicated the presence of LOH, while a value of 1 or more indicated the absence of LOH. Mutations with unknown locus-specific copy numbers in either the CCLE or CLP dataset were determined for LOH status using the other dataset. Samples with mutations that still had unknown LOH were excluded from the relevant analysis.

### Mutational signature 3

Using all SNVs extracted from gene mutation profiles as input, the cosine similarity to the mutational signature 3 of COSMIC version 2 reference signatures was calculated using SigMA (#19).

### HRD score

The scarHRD (#20) was utilized to calculate the telomeric allelic imbalance (TAI) (#11), large-scale state transition (LST) (#12), and genomic loss of heterozygosity (LOH) scores (#13) based on the segmented allele-specific copy number data and the estimated tumor ploidy. The HRD score was determined as the sum of these scores.

### BRCA1 methylation

Gene expression and methylation beta values of the *BRCA1* promoter regions were analyzed to annotate methylation silencing for each cell line. Specifically, in the CCLE dataset, samples with a beta value > 0.3 in two promoter regions located 1Kb upstream from the transcription start site and gene expression < 20% were considered to have *BRCA1* methylation (Supplementary FigureS1A). In the CLP dataset, significant negative correlations were observed between beta values and BRCA1 gene expression in 17 promoter methylation probes. Samples with beta values > 0.2 and gene expression < 30% for at least 15 of the 17 probes were considered to have *BRCA1* methylation (Supplementary FigureS1B).

### Drug sensitivity datasets

Drug sensitivity screening data for GDSC1/2, CTRP1/2, and PRISM were obtained from their websites (Supplementary TableX2). For the gCSI dataset, curated drug response data were obtained from PharmacoDB (#21). We employed the area under the drug-response curve (AUC) values rather than the half maximal inhibitory concentration (IC50) for comparing drug effects, because the AUC reportedly contains fewer outliers compared to the IC50 (#22), and, indeed, had fewer missing values in the datasets. Among the compounds in the CTRP1/2, those with clinical application or FDA approval were retained. Similarly, among the compounds studied in the PRISM, those relevant to the field of oncology were retained. All drugs were integrated using their drug name and PubChem compound ID, and then categorized based on annotations in the GDSC and CTRP. The categories included Platinum, PARP inhibitor, Topoisomerase inhibitor, DNA alkylator, DNA inhibitor, Antimetabolite, PI3K/MTOR signaling, RTK signaling, Chromatin-related, ERK MAPK signaling, Cell cycle-related, Anti-microtubular, Protein stability and degradation, Apoptosis regulation, EGFR signaling, WNT signaling, Genome integrity, Metabolism, Hormone-related, IGF1R signaling, p53 pathway, Cytoskeleton, JNK and p38 signaling, and Others. Assays with insufficient number of samples for comparison were excluded; in the CCLE datasets, 82 assays with fewer than 10 samples with HR-related gene alterations were excluded and the remaining 800 assays were analyzed; in the CLP datasets, 213 assays with fewer than 10 samples with HR-related gene alterations were excluded and the remaining 669 assays were analyzed.

### Responses to Oncology Agents and Dosing in Models to Aid Preclinical Studies (ROADMAPS) dataset (#23)

Binary annotations of drug responsiveness to intraperitoneal (IP) dosing of cisplatin (‘Y’ or ‘N’) were obtained for xenograft models generated by transplanting CCLE cell lines into immunocompromised mice from the published Excel file (roadmaps_12-14-2021.xlsx) in the ROADMAPS project (#23).

### The Breast Cancer PDTX Encyclopaedia (BCaPE) dataset (#24)

Mutational signature 3 was calculated from the published per-sample SNV profiles in the BCaPE (#24) using SigMA (#19).

### Statistical analysis

All statistical analyses were performed using Python (3.8.8). Comparison of drug sensitivity between cell lines per assay was performed by the Mann-Whitney and Spearman’s correlation tests using SciPy module (1.7.2). Differences in continuous values between multiple groups were performed by Kruskal-Wallis test using SciPy module (1.7.2). P values were corrected for multiple testing using the Holm method. Adjusted P value < 0.05 was considered statistically significant.

## Results

### Association between HR-related gene alterations and drug sensitivity

Genomic profiles in the CCLE cell lines and multiple drug response data, including GDSC1/2, CTRP1/2, PRISM, and gCSI, were combined, resulting in a total of 800 assays across 1182 cell lines (Supplementary TableS1, S2). This included 10 assays for platinum agent (Carboplatin 2, Cisplatin 4, Oxaliplatin 4) and 14 assays for PARP inhibitors (Olaparib 6, Talazoparib 3, Veriparib 2, Niraparib 2, Rucaparib 1).

First, we compared the area under the drug-response curve (AUC) of each drug between cell lines with *BRCA1/2* alterations (n=25), which include *BRCA1/2* mutations with locus-specific LOH and *BRCA1* methylations, and those without any HR-related gene mutations (n=872) (Figure1). A higher AUC value of a sample indicates higher resistance to the assay. Among the 10 platinum agent assays, 2 assays (Oxaliplatin 1, Cisplatin 1) showed a trend of higher AUCs in cell lines with *BRCA* alterations (unadjusted P < 0.05, adjusted P ≧0.05), while no assays showed lower AUCs. Among 14 PARP inhibitor assays, none showed a significant difference between the two groups.

**Figure1).**
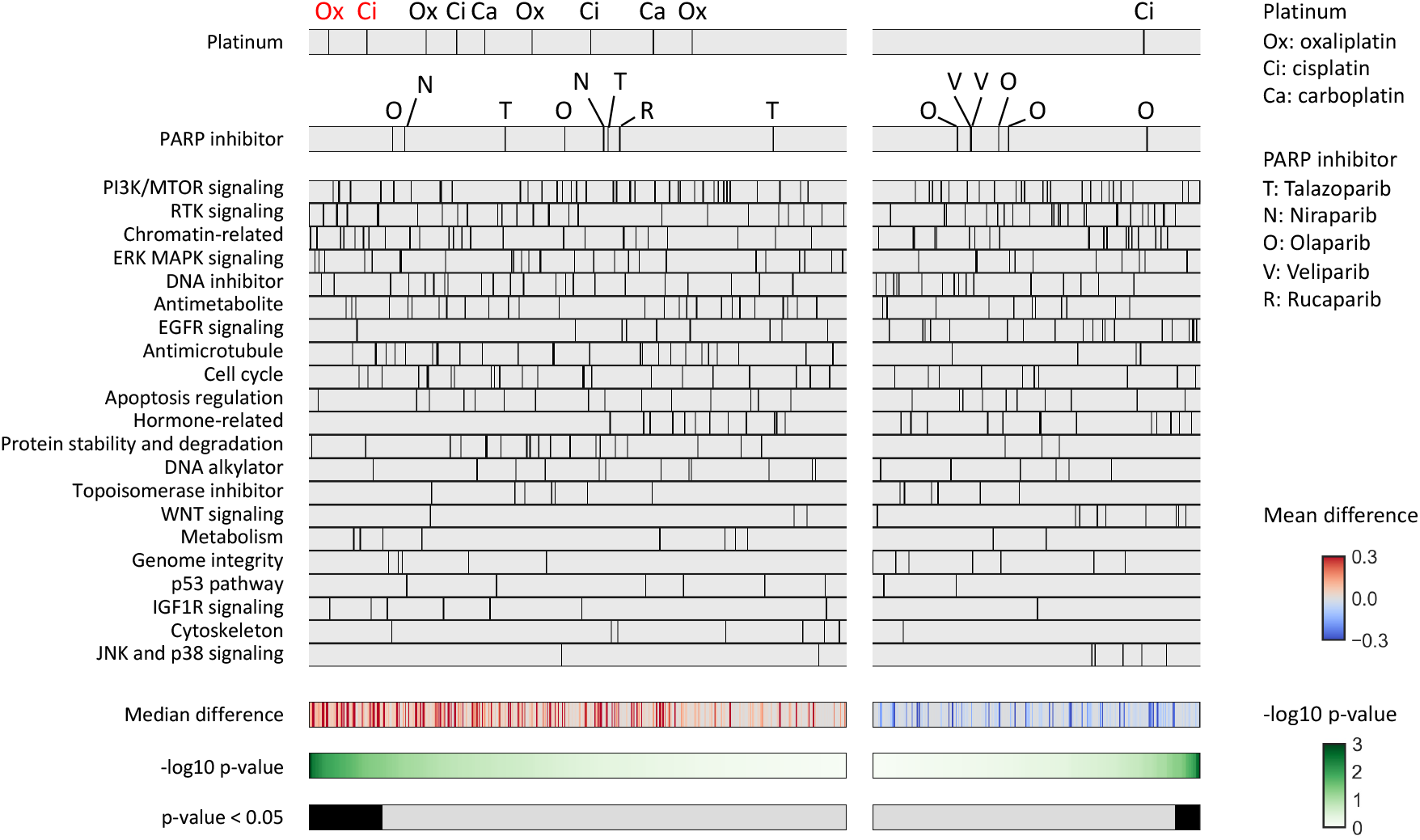
Association between *BRCA1/2* alterations and drug sensitivity. Comparison of the area under the drug-response curve (AUC) between cell lines with *BRCA1/2* alterations (n=25) and no HR-related gene mutations (n=872) was performed. The drugs were divided into left and right panels based on positive and negative median differences, and further ordered based on the lowest and highest p-values (Mann-Whitney test), respectively. Platinum agents (shown in the top panel) and PARP inhibitors (shown in the second panel) exhibiting a trend for positive difference (unadjusted P < 0.05) between the two groups are highlighted in red font.

A similar comparative analysis was performed between cell lines with HR-related gene alterations including *BRCA1/2* (n=82), and those without such alterations (n=872) (Figure2). Two of the 10 platinum agents (Cisplatin 1, Oxaliplatin 1) and 1 of the 14 PARP inhibitors (Niraparib 1) showed a trend of higher AUCs in the HR-related gene-altered samples (unadjusted P < 0.05, adjusted P ≧0.05), and none of them showed lower AUCs.

**Figure2).**
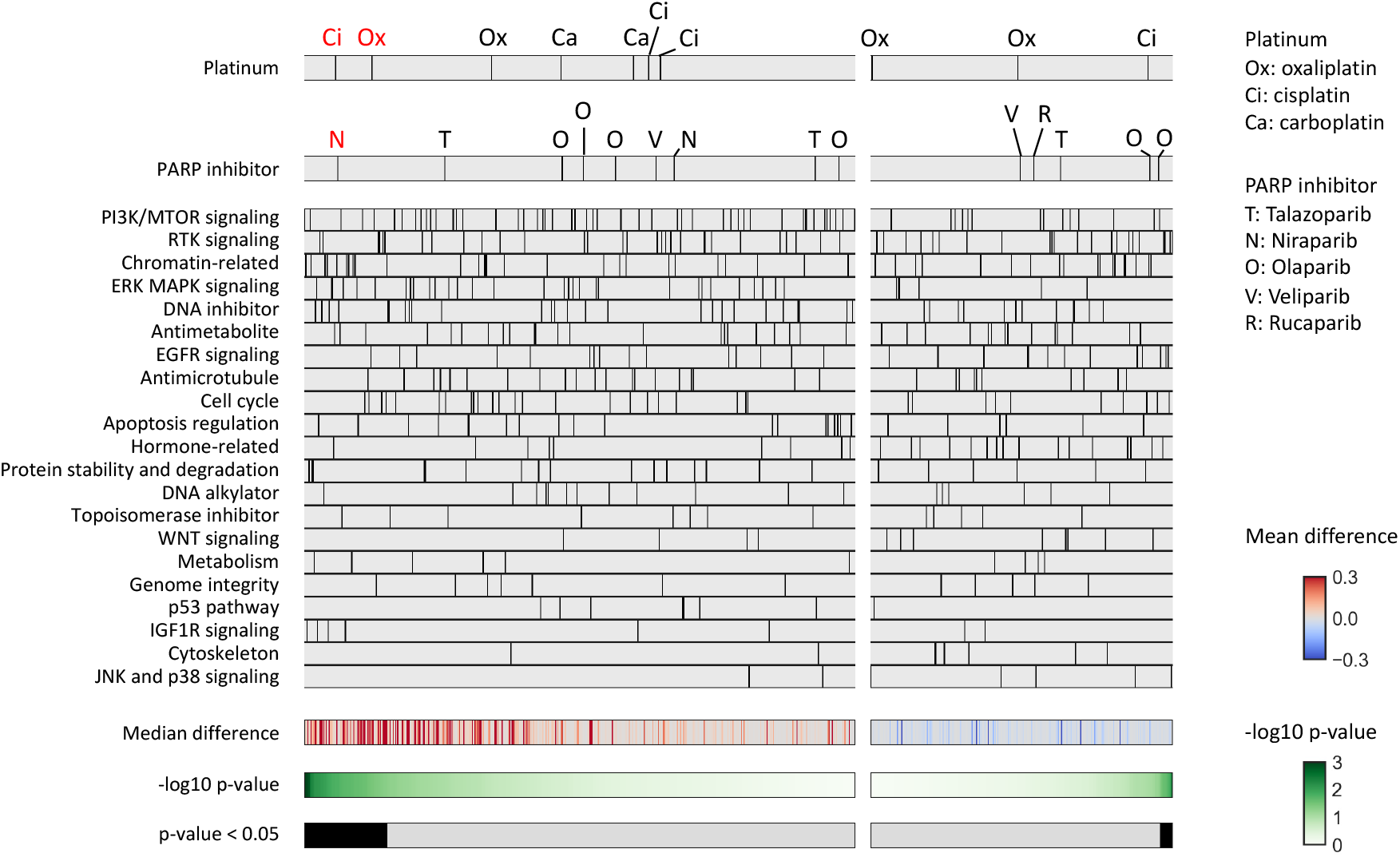
Association between HR-related gene alterations and drug sensitivity. Comparison of the area under the drug-response curve (AUC) between cell lines with HR-related gene alterations including *BRCA1/2* (n=82) and no HR-related gene mutations (n=872) was performed. The drugs were divided into left and right panels based on positive and negative median differences, and further ordered based on the lowest and highest p-values (Mann-Whitney test), respectively. Platinum agents (shown in the top panel) and PARP inhibitors (shown in the second panel) exhibiting a trend for positive differences (Unadjusted P < 0.05) between the two groups are highlighted in red font.

### Genomic scar signatures associated with HRD status in cancer cell lines

Genomic scar signatures, which are indicators of characteristic genomic changes associated with HRD, include the HRD score and the mutational signature 3. The HRD score quantifies the characteristic chromosomal structural changes based on the SNP array data (see methods). The mutational signature 3 is calculated from the pattern of single base substitutions in the context of flanking bases, obtained through genome-wide analysis of somatic mutational profiles. The HRD score and signature 3 were positively correlated in the CCLE dataset (n = 914, Spearman r = 0.475, p = 1.25 ×10^−52^, Figure3A). These two scores differed significantly between groups according to the type of HR-related gene alterations (P = 1.6 ×10^−10^, 2.4 ×10^−7^, respectively, Figure 3B, 3C). They were higher in samples with *BRCA1/2* mutation with LOH, *BRCA1* methylation, and other HR-related gene mutations with LOH compared to those with HR-related gene mutations without LOH or without any HR-related mutations (Figure 3B, 3C). These findings indicate the association between HRD status and genomic scar signatures, commonly observed in clinical samples, was replicated in the cancer cell line dataset.

**Figure3).**
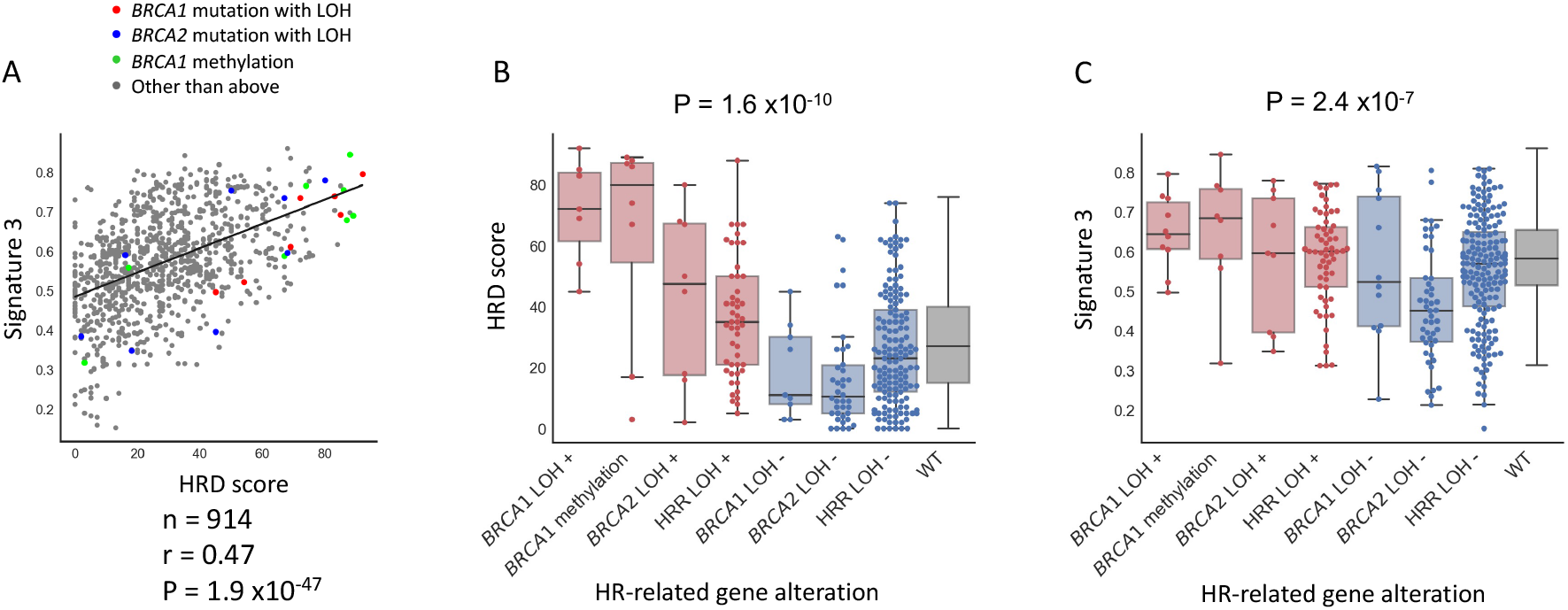
Correlations between HRD status vs genomic scar scores in the CCLE dataset. A) Correlation between HRD score and mutational signature 3 and distribution of BRCA1/2 alterations. HRD scores and mutational signature 3 values were positively correlated. B) Association between HR-related gene alterations with locus-specific LOH and HRD score. HRD scores were significantly different among the groups (Kruskal-Wallis test, P = 1.6 ×10^−10^). Samples with HR-related gene alterations including BRCA1/2 tended to have higher HRD scores. HRR LOH +/-: samples with HR-related gene mutations other than BRCA1/2 with/without the locus-specific LOH. WT; samples without any HR-related mutations. C) Association between HR-related gene alterations with locus-specific LOH and mutational signature 3. Mutational signature 3 were significantly different among the groups (Kruskal-Wallis test, P= 2.4×10^−7^). Samples with HR-related gene alterations including BRCA1/2 tended to have higher signature 3.

### Correlation between genomic scar signatures and drug sensitivity

Spearman’s correlation analysis between HRD scores and AUC (Figure4) showed a trend toward positive correlations in 7 and negative correlations in 1 of the 10 platinum assays (unadjusted P <0.05). After correction for multiple testing, 6 assays (Oxaliplatin 4, Cisplatin 1, and Carboplatin 1) remained statistically significant for positive correlations (adjusted P <0.05). Among the 14 PARP inhibitor assays, 6 (Olaparib 3, Talazoparib 2, Niraparib 1) showed positive correlations both before and after multiple testing corrections (P <0.05), and no assays showed a trend toward negative correlations.

**Figure4).**
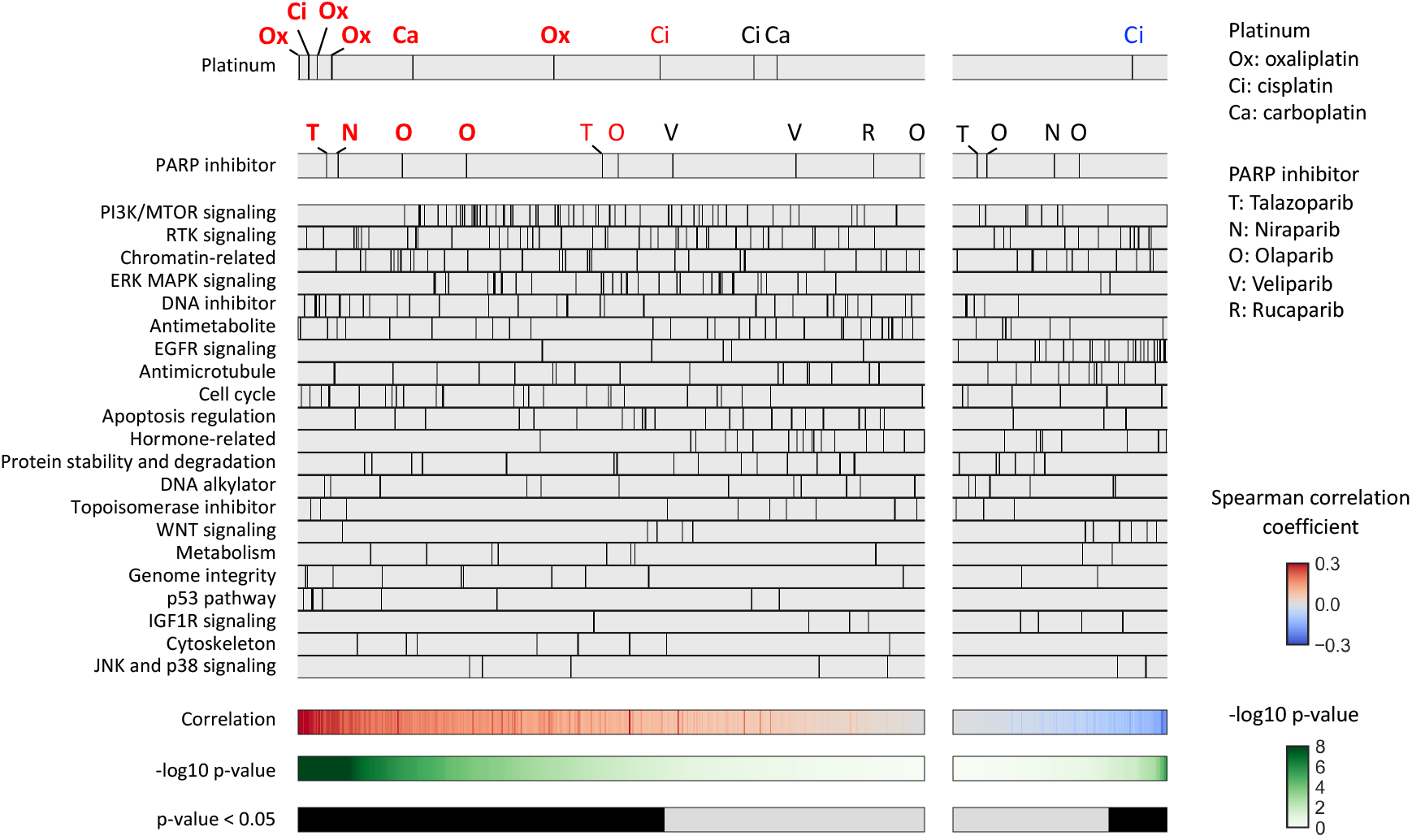
Association between HRD score and drug sensitivity. Spearman’s correlation between HRD score and the area under the drug-response curve (AUC) was analyzed. The drugs were divided into left and right panels based on positive and negative median differences, and further ordered based on the lowest and highest p-values, respectively. Platinum agents (shown in the top panel) and PARP inhibitors (shown in the second panel) exhibiting a trend for positive or negative differences (unadjusted P < 0.05) between the two groups are highlighted in red and blue font, respectively, and those remained statistically significant (adjusted P < 0.05) after multiple testing correction are indicated in bold style.

Similarly, Spearman’s correlation analysis between mutational signature 3 and AUC (Figure5) showed a trend toward positive correlations in 4 of the 10 platinum assays (unadjusted P <0.05) and 2 (Oxaliplatin 2) of them remained statistically significant (adjusted P <0.05). No assays showed significant negative correlations. Among the 14 PARP inhibitor assays, 2 assays showed a trend toward positive correlations (unadjusted P <0.05, adjusted P ≧0.05), and no assays showed a trend toward negative correlations.

**Figure5).**
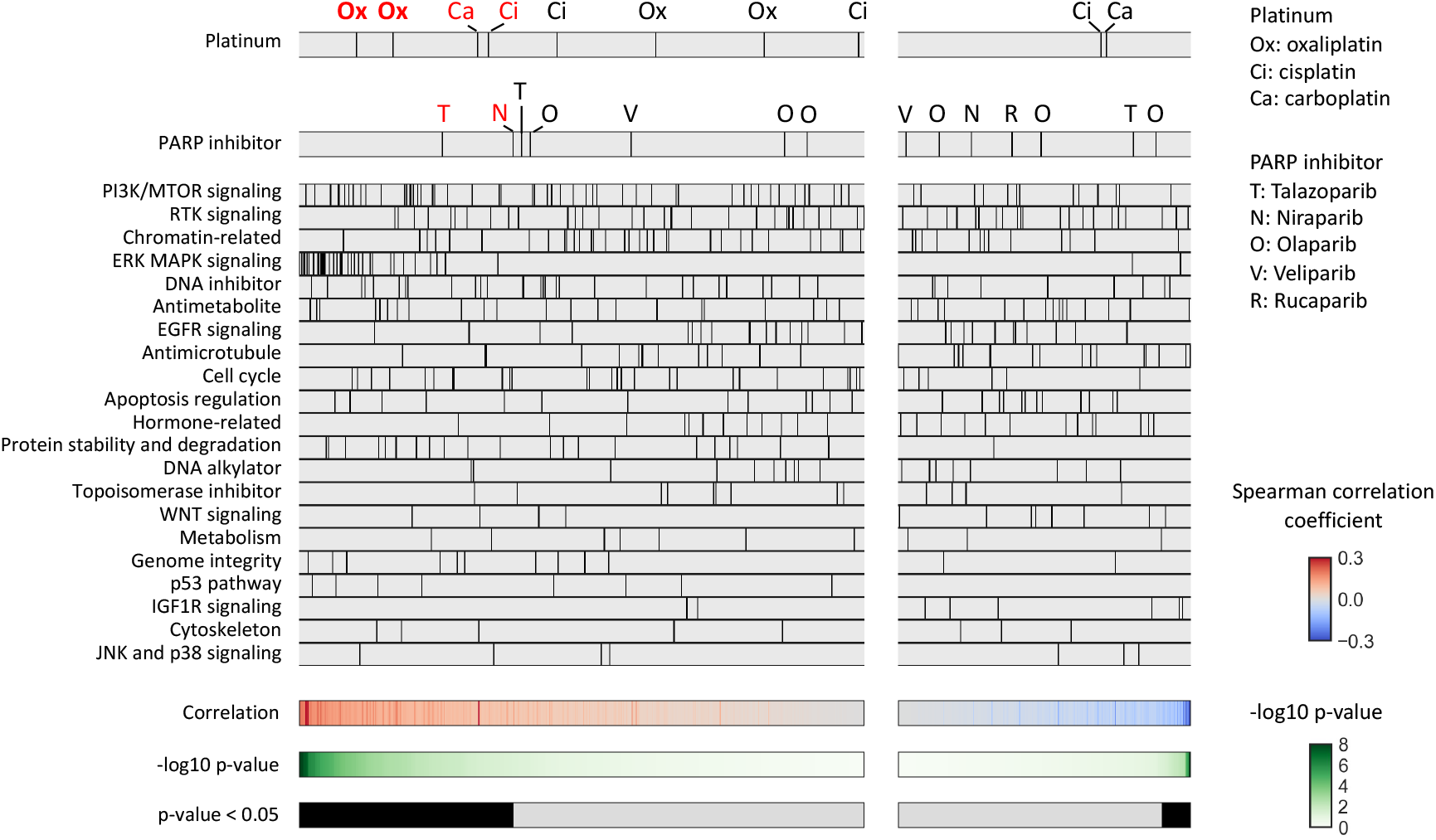
Association between signature 3 and drug sensitivity. Spearman’s correlation between mutational signature 3 value and the area under the drug-response curve (AUC) was analyzed. The drugs were divided into left and right panels based on positive and negative median differences, and further ordered based on the lowest and highest p-values, respectively. Platinum agents (shown in the top panel) and PARP inhibitors (shown in the second panel) exhibiting a trend for positive difference (unadjusted P < 0.05) between the two groups are highlighted in red font, and those remained statistically significant (adjusted P < 0.05) after multiple testing correction are indicated in bold style.

Results were replicated in the same analysis restricted to breast and ovarian cancer cell lines (n = 54 and 62) (Supplementary FigureS2). Furthermore, the same series of analyses using the CLP dataset, in which the molecular profiles were investigated independently of the CCLE, showed that the distributions of HRD scores and mutational signature 3 were similar in the two datasets (Supplementary FigureS3), and the association with drug sensitivity was almost identical (Supplementary FigureS4-S8).

These results indicate that based on analysis of the currently available cancer cell line datasets, HRD status was more likely to correlate with resistance than with sensitivity to platinum or PARP inhibitors, in contrast to data reported in clinical tumors.

### Association between HRD status and drug sensitivity evaluated under different conditions and with different assays from the 2D drug screening assays

All AUC values used in this study were derived from cytotoxic assays, where cell viability was assessed several days after drug administration in a two-dimensional (2D) cell culture setting. We next examined the association between HRD status and drug sensitivity under different assay conditions.

We accessed drug sensitivity data from the ROADMAPS dataset (#23) of cancer cell line-derived xenograft models, which were generated by transplanting CCLE cancer cell lines into immunocompromised mice, and examined the HRD status of the corresponding CCLE cell lines. The HRD score and mutational signature 3 tended to be higher in the cisplatin-sensitive strain compared to the cisplatin-resistant strain (HRD score; median 45 vs. 36, P = 0.41, Mutational signature 3; median 0.63 vs. 0. 56, P = 0.45, Supplementary figure S9A).

In addition, we analyzed mutational profiles of the short-term cultures of ex-vivo tumor cells generated from breast cancer patient-derived tumor xenografts reported by Bruno et.al (#24) and found that the mutational signature 3 and the AUCs of PARP inhibitors tended to be negatively correlated (Olaparib; n = 19, r = -0.43, P = 0.066 Rucaparib; n = 13, r = -0.41, P = 0.17, Supplementary figure S9B) in these datasets.

These results suggest that experimental models in vivo or under different culture conditions than the existing cell line datasets may retain the relationship between HRD status and drug sensitivity observed in clinical tumors.

## Discussion

Many earlier experimental studies have shown that cell lines with HRD are sensitive to platinum and PARP inhibitors even in a two-dimensional (2D) culture setting. Knockout of *BRCA1* or *BRCA2* in human embryonic stem cells or in the chicken B cell-derived cell line DT40 cells markedly increased sensitivity to PARP inhibitors and platinum drugs (#25, #26). During the 2D culture of the pancreatic cancer cell line CAPAN-1 with a *BRCA2* mutation, the addition of cisplatin induced the emergence of subclones with a *BRCA2* reversion mutation, resulting in resistance to cisplatin and the PARP inhibitor AG-14361 (#27). The accumulation of such experimental evidence eventually led to the clinical implementation of personalized treatment with biomarker-based PARP inhibitors today. However, most of these experiments examined the association between dysfunction of BRCA molecules and in vitro drug sensitivity in cells with identical backgrounds. On the other hand, there have been limited reports investigating the association by comparing multiple cell lines derived from different backgrounds. To the best of our knowledge, only one study in 2012 reported a correlation between the TAI score, one of the genomic scar scores for HRD, and sensitivity to cisplatin in 10 breast cancer cell lines (#11). In contrast, a study in 2020 comparing 12 breast cancer cell lines, including *BRCA* mutated lines, reported that *BRCA* status was independent of the response to PARP inhibitors (#28). In the current study, using the largest database of cancer cell lines, we consistently observed that HRD status was not associated with sensitivity to PARP inhibitors or platinum drugs, but rather with resistance.

The discrepancy in drug response between cancer cell lines and clinical tumors observed here can be attributed to several factors. Firstly, clinical tumor tissues comprise not only tumor cells but also mesenchymal cells and immune cells, whose interactions affect tumor growth and drug sensitivity, while cancer cell lines lack such a complex tumor microenvironment. Secondly, the in vitro drug screening assays examined in the cell line databases evaluated drug response over a very short period of 3-5 days, which is considerably shorter than the assessment of efficacy in vivo or in clinical tumors, where it is usually assessed on a weekly or monthly basis. The finding that genomic scar signatures tended to be higher in cisplatin-sensitive strains in the xenograft model (Figure6A) suggests that differences between in vitro and in vivo assays may contribute to the observed discrepancy. Thirdly, the process of establishing immortalized cell lines from tumor tissues inevitably involves significant selection pressure. During this process, fast-proliferating clones may dominate the heterogeneous cell population of a tumor, or clones that have acquired advantageous traits for growth on 2D plates through gene mutations during cell culture may be preferentially selected (#29). It has been reported that disruption of the HR pathway in cells, such as through knockout of the *BRCA1/2* genes, leads to suppression of cell proliferation (#30, #31). This is likely because cells deficient in HR function use alternative DNA repair pathways to repair DNA damage while the cell cycle checkpoint is activated and proliferation is halted. However, given intratumor heterogeneity, when attempting to establish cancer cell lines from tumors with HRD, slow-growing cells may be eliminated and fast-growing cells with highly enhanced alternative mechanisms may be selected.

Consequently, the established cell line library may possess different properties compared to clinical tumors. The trend that HRD status correlates with sensitivity to PARP inhibitors in short term in vitro culture of PDX-derived tumor cells in breast cancer (Figure6B) provides support to this hypothesis.

The collective findings of this study indicate that PDX models offer a superior approach compared to 2D culture of cancer cell lines for investigating clinically relevant drug effects in preclinical studies. It has long been well documented that in vivo drug studies using PDX models more closely resemble clinical tumors than in vitro studies with cancer cell lines (#32, #33). Specifically, in the context of HRD and PARP inhibitors, the tumor shrinkage effect of niraparib treatment on ovarian cancer PDX models was greater in tumors with *BRCA* mutations and higher HRD scores, as reported by the pharmaceutical company in the application form to the Pharmaceuticals and Medical Devices Agency in Japan. (https://www.pmda.go.jp/files/000245811.pdf, Table 8). The National Cancer Institute has performed comprehensive molecular characterization of a large number of human tumors, their corresponding PDX models, and subsequent passages in various types of cancer using next-generation sequencing, and the results have been published in the Patient-Derived Models Repository (PDMR) database (#34). The addition of drug sensitivity data to these resources holds significant promise for advancing cancer drug research in the future. In addition, 3D in vitro cultures such as patient-derived organoids have been utilized in recent studies because they behave more like clinical tumors than 2D cell lines and can be cultivated with higher efficiency than PDX models (#35, #36). However, it has been reported that even in organoids, drug sensitivity may or may not accurately reflect that of clinical tumors, depending on the culture medium and the type of drug (#37). Therefore, careful consideration of experimental conditions is required to ensure accurate evaluation of drug responses in organoids.

In conclusion, we analyzed molecular characteristics and drug screening data from the largest currently available cancer cell line library and found that differences in HRD status between cell lines do not correlate with sensitivity to platinum or PARP inhibitors in 2D cell culture assays. This critical deviation from the anticipated findings in clinical oncology may also apply to molecularly-targeted drugs other than PARP inhibitors, warranting caution among researchers using the database for future cancer research. The findings in this study emphasize the importance of utilizing more advanced models, including patient-derived xenografts and organoids, to conduct highly beneficial preclinical studies that can improve personalized medicine for cancer patients.

## Acknowledgements

The authors thank Hillman Robert Tyler for his helpful discussions and support in editing the manuscript.

## Authors contributions

ST: Conceptualization, Software, Formal analysis, Writing - Original Draft, Visualization

KM: Validation, Resources, Data Curation, Supervision,

NM: Conceptualization, Methodology, Investigation, Data Curation, Writing - Original Draft

All authors; Writing - Review & Editing

## Data and code availability

Data sources of cancer cell line datasets are summarized in Supplementary TableX2. The processed data and codes to reproduce the results of this work are available on the GitHub project page (https://github.com/shirotak/CellLine_HRD_DrugRes). Other codes for data preprocessing are available from the corresponding author upon reasonable request.

## Ethics

This study used only previously published anonymized data and was exempt from institutional review board approvals and informed consent for patients.

