## Supplementary data for "Homologous recombination deficiency in cell line libraries does not correlate with in vitro sensitivity to platinum agents or PARP inhibitors"

#### Supplementary Figures

**Figure S1) Annotation of epigenetic silencing of BRCA1 promoter methylation**

**Figure S2) Analyses in breast (n=54) and ovarian (n=62) cell lines in the CCLE dataset**

**Figure S3) The distributions of HRD scores and mutational signature 3 between CCLE and CLP datasets**

**Figure S4) Association between BRCA1/2 alterations and drug sensitivity in the CLP dataset**

**Figure S5) Association between HR-related gene alterations and drug sensitivity in the CLP dataset**

**Figure S6) Correlations between HRD status vs genomic scar scores in the CLP dataset**

**Figure S7) Association between HRD score and drug sensitivity in the CLP dataset**

**Figure S8) Association between signature 3 and drug sensitivity in the CLP dataset**

**Figure S9) Association of HRD status and drug sensitivity in assay conditions different from that of the cell line databases.**

#### Supplementary Tables

**Table X1) Sources for molecular profiling of cancer cell lines**

**Table X2) Sources of drug response data**

**Figure S1) Annotation of epigenetic silencing of BRCA1 promoter methylation**

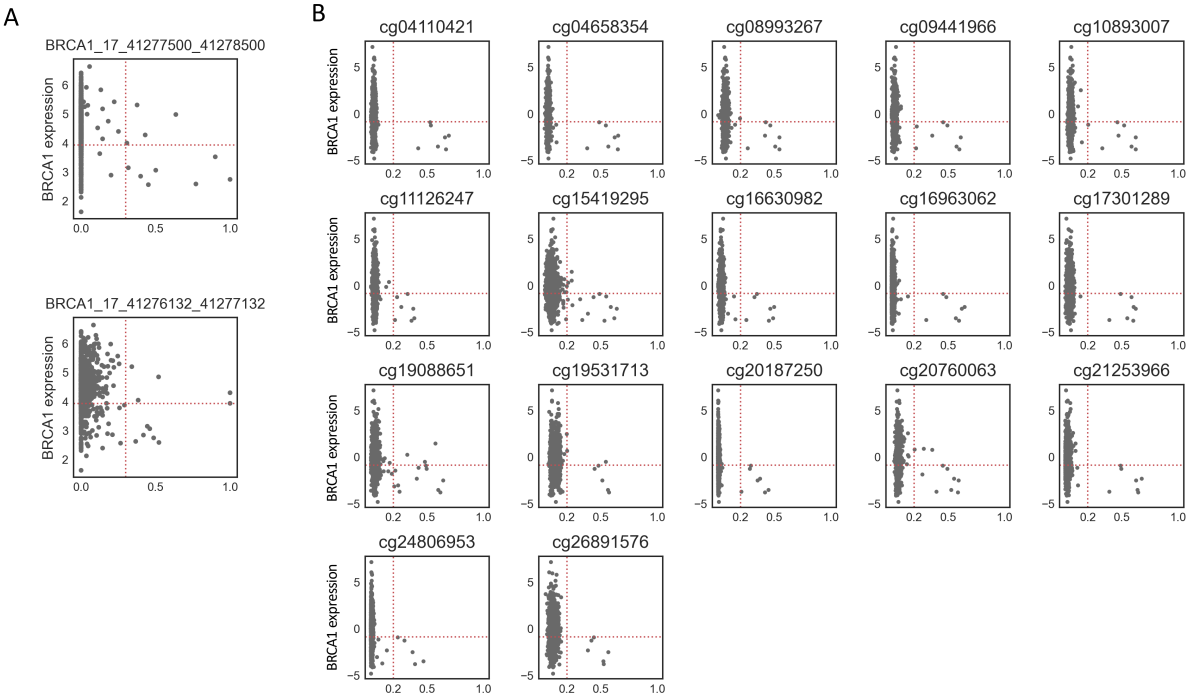


**A) CCLE dataset**

Methylation beta values (X-axis) of the two *BRCA1* promoter regions (located 1Kb upstream from the transcription start site) and gene expression (Y-axis) were plotted. Samples with a beta value > 0.3 in two promoter regions and gene expression < 20% were considered to have *BRCA1* methylation.

B) **CLP dataset**

Methylation beta values (X-axis) and BRCA1 gene expression (Y-axis) were significant negatively correlated in 17 promoter methylation probes. Samples with beta values > 0.2 and gene expression < 30% for at least 15 of the 17 probes were considered to have BRCA1 methylation.

**Figure S2) Analyses in breast (n=54) and ovarian (n=62) cell lines in the CCLE dataset**

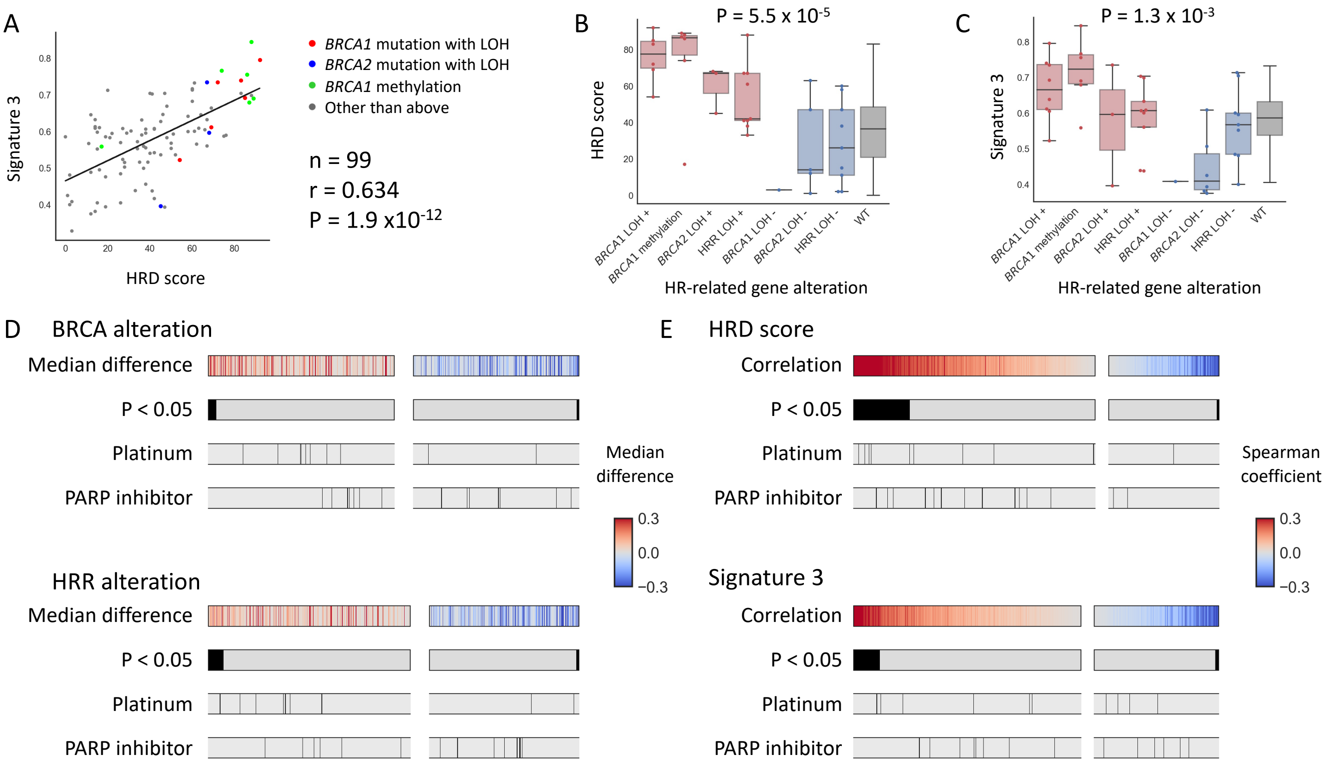


A) Correlation between HRD score and mutational signature 3 and distribution of BRCA1/2 alterations
HRD scores and mutational signature 3 values were positively correlated.

B) Association between HR-related gene alterations with locus-specific LOH and HRD score.
HRD scores were significantly different among the groups (Kruskal-Wallis test, P = 5.5 x10$^{-5}$).
HRR LOH +/-: samples with HR-related gene mutations other than BRCA1/2 with/without the locus-specific LOH. WT; samples without any HR-related mutations.

C) Association between HR-related gene alterations with locus-specific LOH and mutational signature 3.
Signature 3 were significantly different among the groups (Kruskal-Wallis test, P = 1.3 x10$^{-3}$)

D) Association between drug sensitivity and BRCA1/2 alterations (upper) or HR-related gene alterations.
Comparison of the area under the drug-response curve (AUC) between cell lines with BRCA1/2 alterations (n=16) or HR-related gene alterations (n=25) vs no HR-related gene mutations (n=83) was performed. The drugs were divided into left and right panels based on positive and negative median differences, and further ordered based on the lowest and highest p-values (Mann-Whitney test), respectively.

E) Spearman's correlation analysis between the drug-response curve (AUC) vs HRD score (upper) or signature 3 (lower).
The drugs were divided into left and right panels based on positive and negative median differences, and further ordered based on the lowest and highest p-values, respectively.

**Figure S3) The distributions of HRD scores and mutational signature 3 between CCLE and CLP datasets**

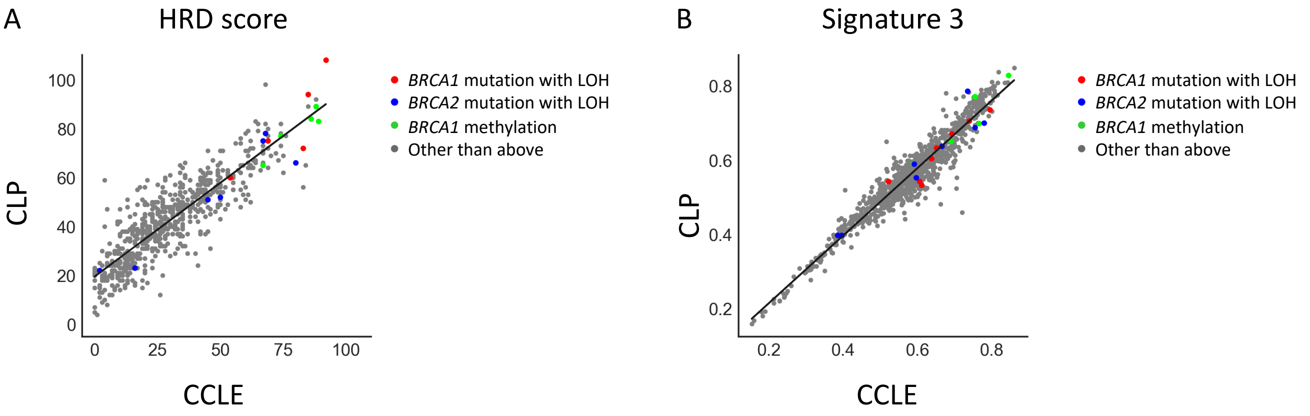


n = 636
rS = 0.87
p = 9.8 $x10^{-197}$

n = 941
rS = 0.96
p < 1.0 $x10^{-199}$

The distribution of HRD scores (A) and mutational signature 3 values (B). Identical cell lines in the two data sets showed similar values to each other.

**Figure S4) Association between BRCA1/2 alterations and drug sensitivity in the CLP dataset**

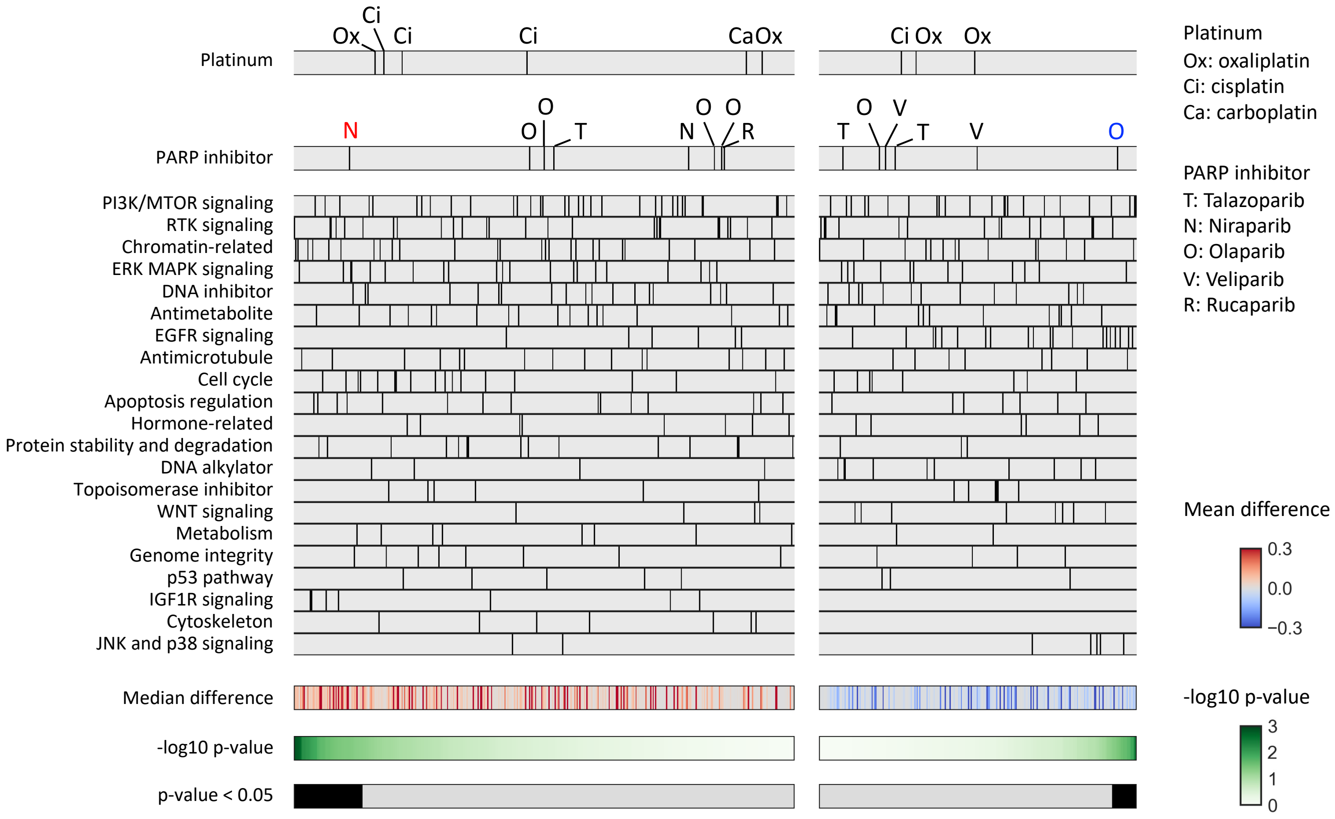


Comparison of the area under the drug-response curve (AUC) between cell lines with BRCA1/2 alterations (n=18) and no HR-related gene mutations (n=774) was performed for each of a total of 669 assays.

The drugs were divided into left and right panels based on positive and negative median differences, and further ordered based on the lowest and highest p-values (Mann-Whitney test), respectively.

Platinum agents (shown in the top panel) and PARP inhibitors (shown in the second panel) exhibiting positive or negative differences (unadjusted P value < 0.05) between the two groups are highlighted in red and blue font, respectively.

**Figure S5) Association between HR-related gene alterations and drug sensitivity in the CLP dataset**

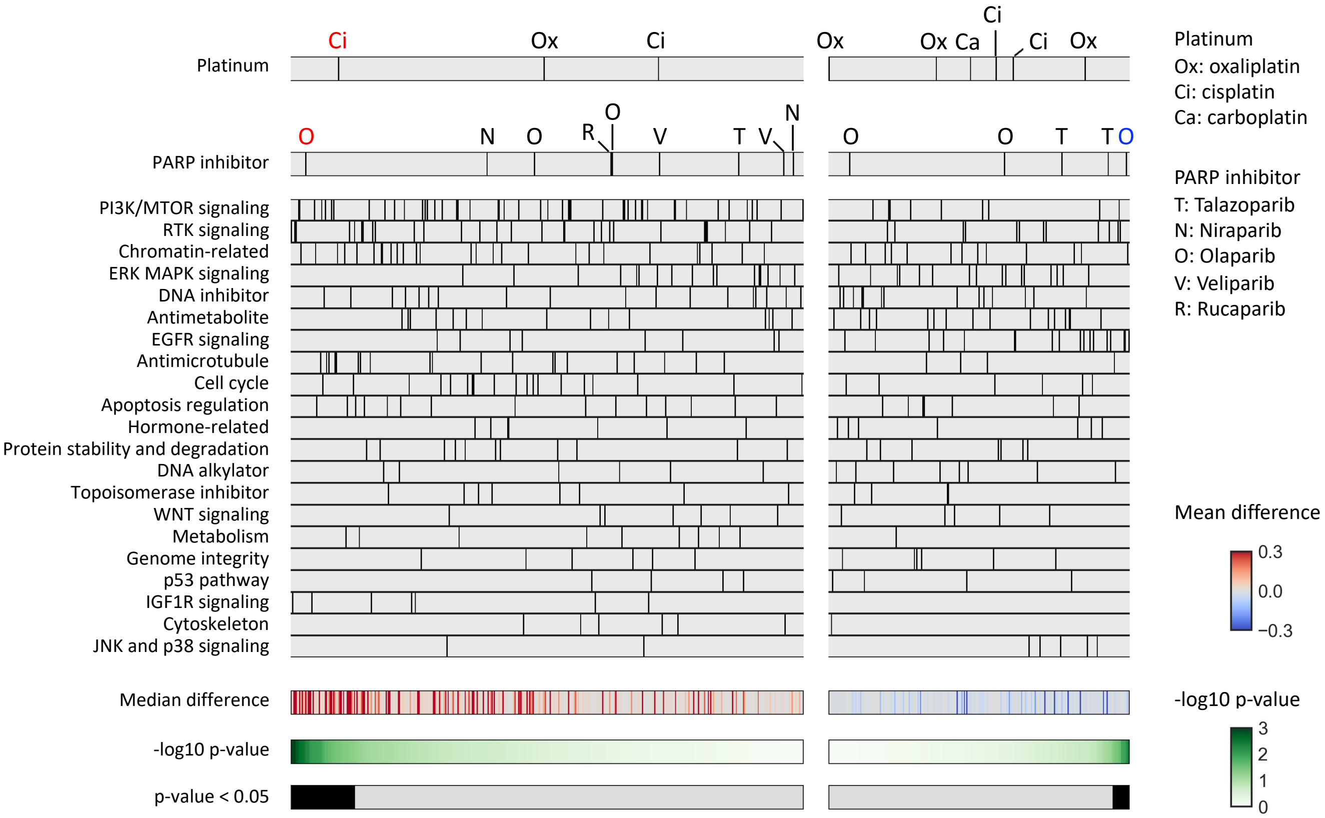


Comparison of the area under the drug-response curve (AUC) between cell lines with HR-related gene alterations, including BRCA1/2, (n=50) and no HR-related gene mutations (n=774) was performed for each of a total of 669 assays.
The drugs were divided into left and right panels based on positive and negative median differences, and further ordered based on the lowest and highest p-values (Mann-Whitney test), respectively.
Platinum agents (shown in the top panel) and PARP inhibitors (shown in the second panel) exhibiting positive or negative differences (unadjusted P value) between the two groups are highlighted in red and blue font, respectively.

**Figure S6) Correlations between HRD status vs genomic scar scores in the CLP dataset**

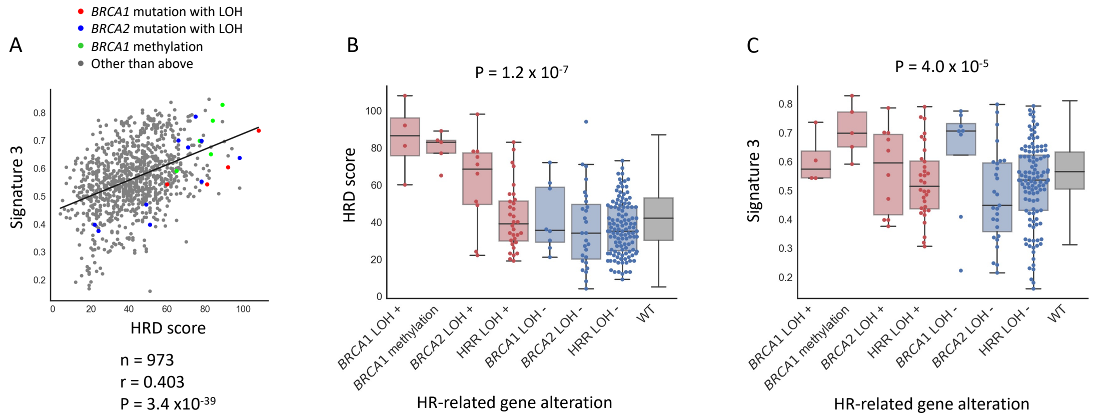


A) Correlation between HRD score and mutational signature 3 and distribution of BRCA1/2 alterations
HRD scores and mutational signature 3 values were positively correlated.

B) Association between HR-related gene alterations with locus-specific LOH and HRD score.
HRD scores were significantly different among the groups (Kruskal-Wallis test, $P = 1.2 \times 10^{-7}$)

C) Association between HR-related gene alterations with locus-specific LOH and mutational signature 3.
Mutational signature 3 were significantly different among the groups (Kruskal-Wallis test, $P = 4.0 \times 10^{-5}$)

**Figure S7) Association between HRD score and drug sensitivity in the CLP dataset**

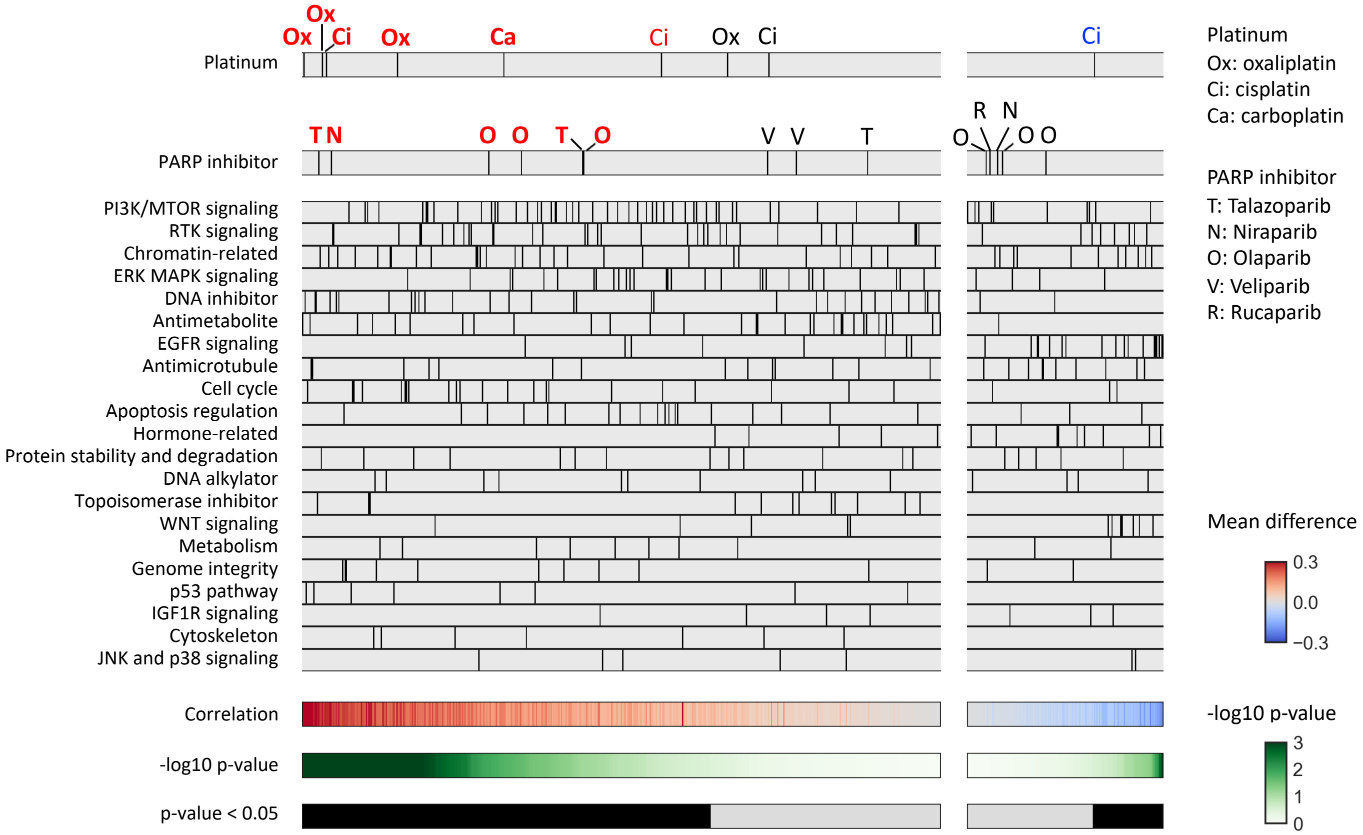


Spearman's correlation between HRD score and the area under the drug-response curve (AUC) was analyzed for each of a total of 669 assays.

The drugs were divided into left and right panels based on positive and negative median differences, and further ordered based on the lowest and highest p-values, respectively. Platinum agents (shown in the top panel) and PARP inhibitors (shown in the second panel) exhibiting positive or negative differences (unadjusted P value < 0.05) between the two groups are highlighted in red or blue font, respectively, and those remained statistically significant (adjusted P value < 0.05) after multiple testing correction are indicated in bold style.

**Figure S8) Association between signature 3 and drug sensitivity in the CLP dataset**

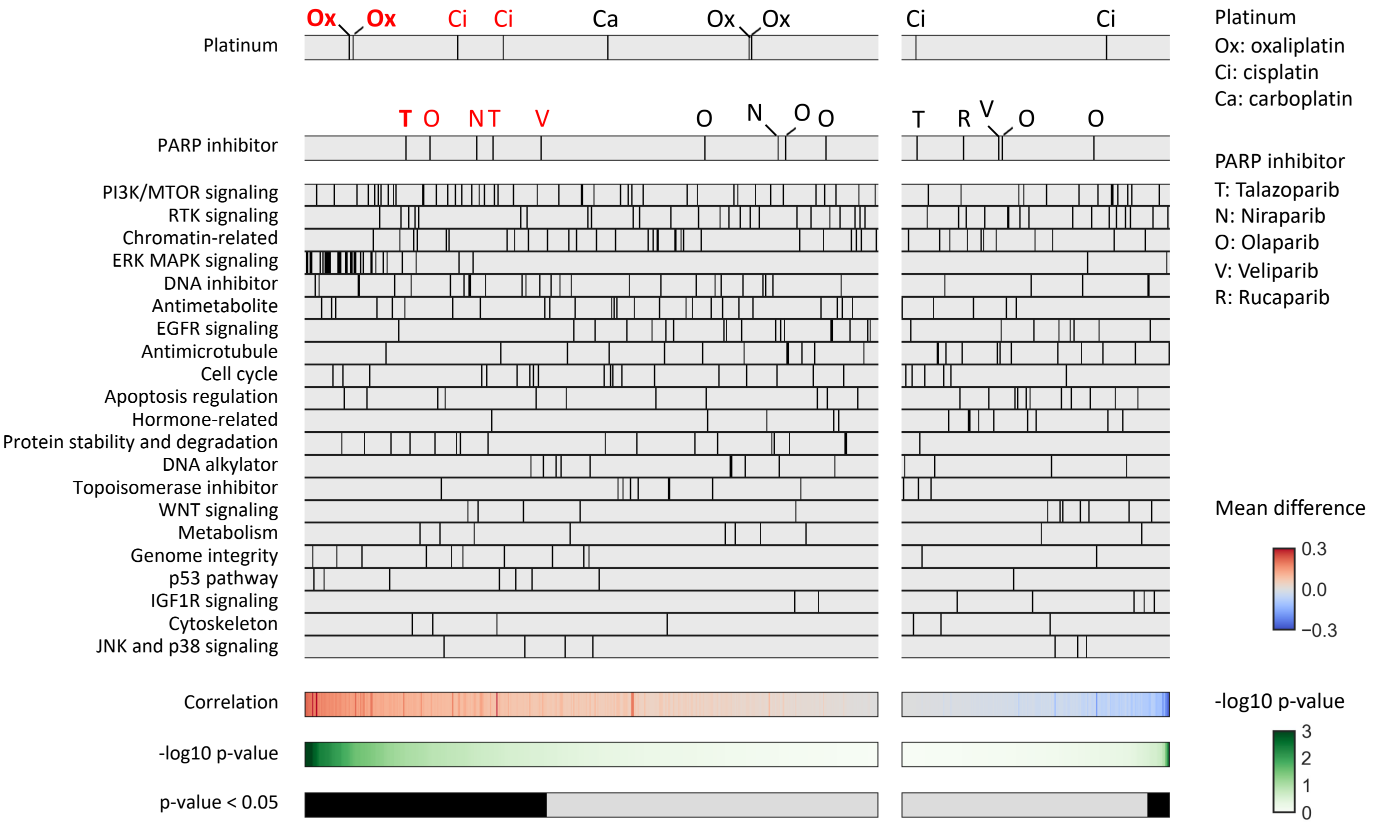


Spearman's correlation between mutational signature 3 value and the area under the drug-response curve (AUC) was analyzed for each of a total of 669 assays.
The drugs were divided into left and right panels based on positive and negative median differences, and further ordered based on the lowest and highest p-values, respectively.
Platinum agents (shown in the top panel) and PARP inhibitors (shown in the second panel) exhibiting positive or negative differences (unadjusted P value < 0.05) between the two groups are highlighted in red or blue font, respectively, and those remained statistically significant (adjusted P value < 0.05) after multiple testing correction are indicated in bold style.

**Figure S9) Association of HRD status and drug sensitivity in assay conditions different from that of the cell line databases.**

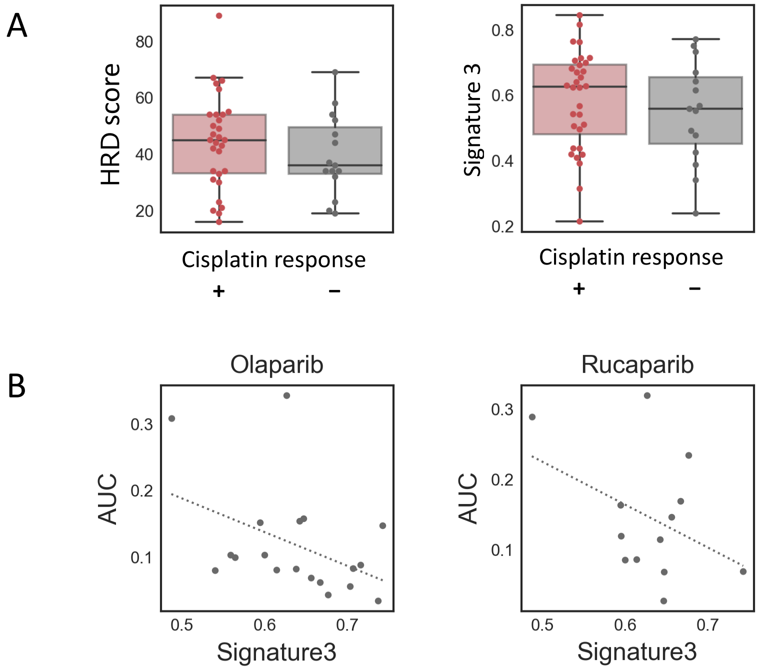


A) Association between cisplatin responsiveness of mouse xenografts implanted with cancer cell lines from the CCLE and the HRD score (left) or signature 3 (right), as analyzed in the ROADMAP dataset.
HRD scored and mutational signature 3 values tended to be higher in the cisplatin-sensitive strain compared to the cisplatin-resistant strain. (HRD score; median 45 vs. 36, p= 0.41, Mutational signature 3; median 0.63 vs. 0. 56, p=0.45, Mann-Whitney test).

B) Association of mutational signature 3 and PARP inhibitor sensitivity in a short-term culture system of ex-vivo tumor cells generated from breast cancer patient-derived tumor xenografts, as reported by Bruna et al.
Mutational signature 3 values and the area under drug response curve (AUC) tended to be negatively correlated in Olaparib (left) and Rucaparib (right) treatment. (Olaparib; r=-0.43, p=0.066 Rucaparib; r=0.41, p=0.17, Spearman' correlation).

**Table X1) Sources for molecular profiling of cancer cell lines**

| | **Sanger, COSMIC-CLP** | **Broad, CCLE** |
|---|---|---|
| Data type | URL, filename | |
| Mutation profiles | https://cancer.sanger.ac.uk/cell_lines/download<br>CosmicCLP_MutantExport.tsv.gz | https://depmap.org/portal/<br>DepMap Public 22Q2, CCLE_mutations.csv |
| Copy number variations | https://cancer.sanger.ac.uk/cell_lines/download<br>cell_lines_copy_number.csv | https://depmap.org/portal/<br>CCLE2019,CCLE_ABSOLUTE_combined_20181227.xlsx |
| | https://cancer.sanger.ac.uk/cell_lines/download<br>PICNIC_average_ploidies.tsv | https://depmap.org/portal/download/custom/<br>Aneuploidy.csv |
| Gene expression | https://cancer.sanger.ac.uk/cell_lines/download<br>CosmicCLP_RawGeneExpression.tsv.gz | https://depmap.org/portal/download/custom/<br>Expression_22Q2_Public.csv |
| DNA methylation | https://www.ncbi.nlm.nih.gov/geo/query/acc.cgi?acc=GSE68379<br>GSE68379_Matrix.processed.txt.gz | https://depmap.org/portal/download/custom/<br>Methylation_(1kb_upstream_TSS).csv |

**Table X2) Sources of drug response data**

| | URL, Source name | Drug-response AUC | Drug annotation | Nr of drug x cell | Measurement (assay) |
|---|---|---|---|---|---|
| GDSC1 | https://www.cancerrxgene.org/downloads/bulk_download GDSC1-dataset | GDSC1_fitted_dose_response_25Feb20.xlsx AUC | screened_compounds_rel_8.4.csv, TARGET_PATHWAY | 240 X 987 | 72h drug treatment fluorescence-based assay |
| GDSC2 | https://www.cancerrxgene.org/downloads/bulk_download GDSC2-dataset | GDSC2_fitted_dose_response_25Feb20.csv AUC | | 166 X 809 | |
| CTRP1 | https://ctd2-data.nci.nih.gov/Public/Broad/CTRPv1.0_2013_pub_Cell_154_1151/ | v10.D3.area_under_conc_curve.txt area_under_curve | CTRPv1.0._INFORMER_SET.xlsx, target_or_activity_of_compound & cpd_status # | 90 X 240 | 72h drug treatment CellTiter-Glo assay |
| CTRP2 | https://ctd2-data.nci.nih.gov/Public/Broad/CTRPv2.0_2015_ctd2_ExpandedDataset/ | v20.data.curves_post_qc.txt area_under_curve | CTRPv2.0._INFORMER_SET.xlsx, target_or_activity_of_compound & cpd_status # | 173 X 887 | |
| PRISM | https://depmap.org/portal/download/ PRISM Repurposing 19Q4 | secondary-screen-dose-response-curve-parameters.csv auc | secondary-screen-dose-response-curve-parameters.csv moa & disease.area $ | 172 X 481 | PRISM assay https://www.theprismlab.org/the-prism-assay/ |
| gCSI | https://pharmacodb.pmgenomics.ca/pharmacogx?pgx=4 gCSI (from PharmacoDB) | gCSI (R/PharmacoGx) GR_AOC_published | gCSI (R/PharmacoGx) (drug name) | 41 X 569 | 72h drug treatment CellTiter-Glo assay. |

### From “cpd_status” annotation, those with “clinical, FDA approved” were retained

$ From “disease.area” annotation. those with relation to “oncology, malignancy” were retained
